# REPRODUCTIVE COMPATIBILITY OF TWO LINES OF *DELIA PLATURA* (DIPTERA: ANTHOMYIIDAE)

**DOI:** 10.1101/2023.08.01.551352

**Authors:** Allen Bush-Beaupré, Marc Bélisle, Anne-Marie Fortier, François Fournier, Andrew MacDonald, Jade Savage

**Affiliations:** Université de Sherbrooke, Département de biologie, Sherbrooke, Canada; Bishop’s University, Department of biology, Canada; Centre d’étude de la forêt (CÉF); Compagnie de recherche Phytodata inc, Sherrington, Canada; Collège Montmorency, Laval, Canada

**Author notes:** Corresponding author: Allen Bush-Beaupré, Université de Sherbrooke, Département de biologie, 2500 Boulevard de l’Université, Sherbrooke, Québec, Canada, J1K 2R1.

## Abstract

Accurate identification of agricultural pests is a major component of integrated pest management. The seedcorn maggot, *Delia platura* (Diptera: Anthomyiidae), is a cosmopolitan polyphagous pest species which may be found in high numbers in numerous crops. Two morphologically identical genetic lines of *D. platura* (H- and N- lines) with distinct distributions were recently identified. However, no study to date has investigated the reproductive compatibility of the two lines and thus the possibility that they may actually be two unique biological entities. A previous study described the reproductive traits of the two lines and suggested that H-line females are highly selective towards the male with which they mate, pointing to a possible pre-mating isolation mechanism between the lines. Using laboratory-reared colonies originating from the Montérégie region in Québec, this study investigates the reproductive compatibility of the two *D. platura* lines. We found that only one of thirty H-line females was inseminated by an N-line male, further suggesting mate choice as a pre-mating isolation mechanism between the lines. However, N-line females were readily inseminated by H-line males, suggesting a lack of pre-mating isolation in this type of cross. The eggs laid by N-line females mated with H-line males had a lower hatching rate than the ones laid by females of intra-line crosses suggesting either post-mating pre-zygotic or post-zygotic partial isolation. However, the larvae that did hatch had a comparable developmental success to those from intra-line crosses in terms of survival and developmental time from larval hatching to adult emergence, pupal weight, and adult sex ratio, suggesting a lack of post-zygotic isolation for these life stages. Considering the different biological traits of the two lines, we suggest the use of the ‘biotype’ terminology to designate the two biological entities and discuss their implications for integrated pest management.

## Introduction

Over the years, the concept of a species has been periodically redefined (Darwin 1859; Dobzhansky 1937; Mayr 1942; Van Valen 1976). Mayden (1997) reported 24 variations of the definition, displaying the transformation and, to a certain point, subjectivity of the ‘species’ concept and terminology. Special care must therefore be applied to avoid following a specific concept, which delimitates species using standards intrinsic to its own philosophy, often conflicting with other concepts. While the conceptualization of what defines a species is relevant to fundamental or taxonomical issues, the study of diverged biological traits and degree of reproductive isolation between populations can also have a bearing in some applied contexts. For instance, subpopulations of a named insect pest species may differ in their biological traits, having major implications in their management (Gomi et al. 2003). Regardless of the precise definition of what a species actually is, delimitation analyses between populations are strongest when assessing both genetic and biological lines of evidence and their congruence (Carstens et al. 2013). During the process of species delimitation, biological differences between populations may be uncovered and, while taxonomical status may yet be determined, terminologies such as ‘biotype’ may be designated to such populations to emphasize their biological differences (Diehl and Bush 1984). Such designation may serve the agronomic community and guide future research on control methods tailored to each biotype.

Following divergence, populations may have evolved certain traits linked to reproduction such that inter-breeding is impeded when they come into contact. These barriers to gene flow are apparent through pre-mating, post-mating pre-zygotic, and post-zygotic isolation factors (Coyne and Orr 2004). Multiple forms of reproductive isolation may thus occur between two populations, with weak effects when observed individually but resulting in a stronger isolation effect when combined (Matsubayashi and Katakura 2009). However, this is not always the case; apparent gene flow barriers do not necessarily amount to complete isolation (Nosil 2007).

Pre-mating isolation between two populations can involve a wide variety of mechanisms. For two individuals to mate, they must recognize one another as being part of compatible populations. The sensory systems involved in mate recognition include the visual (Fordyce et al. 2002), auditory (Boumans & Johnsen, 2014), and chemosensory (Ferveur, 2005; Smadja & Butlin, 2009; Bengtsson et al., 2014) systems and are major components of pre-mating isolation. Other mechanisms of pre-mating isolation include allochronic isolation, where the timing of life history traits differs between populations such as different timing of adult emergence from puparia (Hippee et al. 2016) or mating times (Miyatake et al. 2002). Mechanical incompatibilities due to differences in genital morphology may also contribute to pre-mating isolation (Sánchez-Guillén et al. 2012; Nava-Bolaños et al. 2017).

In the event that copulation between members of two populations occurs, hybridization is not an obligate result. Post-mating, pre-zygotic isolation can occur through differences in internal genital morphology between heterospecific sexes, either through mechanical incompatibility, interactions with sensory mechanisms (Masly 2012), or through inappropriate behavior by either sex during copulation which may lead to sperm transfer failure (Buellesbach et al. 2014). While post-mating pre-zygotic isolation is still a developing area of research, many studies have also demonstrated the involvement of seminal fluid proteins in egg fertilization failure and defective sperm storage (Marshall et al. 2009; Larson et al. 2012; Ahmed-Braimah 2016; Garlovsky et al. 2020; Hill et al. 2021).

The formation of a hybrid is still not a guarantee for reproductive compatibility between populations. Post-zygotic isolation mechanisms include reduced hybrid viability where hybrids may differ from their parental populations in terms of decreased survival (dos Santos et al. 2001), differential developmental time (Ording et al. 2010) and offspring weight (Nakakita and Imura 1981). Additionally, Haldane’s rule predicts that hybrids of the heterogametic sex will typically suffer from reduced performance in terms of development through various mechanisms (reviewed in Laurie, 1997). As such, hybrid offspring sex ratio may be biased towards the homogametic sex (Gibson et al. 2013; Phadnis et al. 2015). Post-zygotic isolation also manifests in the form of hybrid suppression where hybrids may succumb to disruptive sexual selection through strong assortative mating (Naisbit et al. 2001; Tadeo et al. 2018). The genetic inheritance pattern of host preference and performance may also cause hybrids to prefer laying eggs on hosts on which the larvae will experience reduced survival (Forister 2005). Thus, multiple mechanisms may impede hybridization between populations.

While isolating mechanisms between populations are both diverse and numerous, they can also be incomplete, allowing gene flow to occur under certain circumstances. In the animal kingdom, hybridization between insects is the most frequently observed (Schwenk et al. 2008). Such hybridization may facilitate colonization in new environments (Andersen et al. 2019) and/or increase resistance to pest management methods (Anderson et al. 2018), two processes that may result in new or more intense agricultural pest outbreaks (Corrêa et al. 2019). It is thus crucial to investigate hybridization potential between populations and monitor the ensuing hybrids in the field to develop control methods tailored to the targeted population.

While synthetic pesticides are often employed to manage pest populations, their toxicity to non-target organisms (DeLorenzo et al. 2001; Wigle et al. 2009), the environment (Ritter 1990; Taiwo 2019) and human health (Perry 2008; Wigle et al. 2009) is well known. There is thus a need to develop alternative methods to control insect pests. These methods, such as the sterile insect technique (Pereira et al. 2007; Ahmad et al. 2018) or species-specific biological control agents (Davidson and Chandler 2005; Jensen et al. 2006; Eilenberg and Jensen 2018) are specific to the target species and thus rely on accurate identification of the pest in question.

The seedcorn maggot, *Delia platura*, (Meigen) (Diptera: Anthomyiidae) is an agricultural pest with a high diversity of larval hosts including commonly cultivated vegetables and field crops (Hough-Goldstein and Hess 1984; Griffiths 1993; Howard et al. 1994; Soroka and Dosdall 2011). Two morphologically identical genetic clusters (H- and N-line) have been identified within *D. platura* (Savage et al. 2016). The two clusters are separated by a minimum p-distance of 4.45% for the barcoding gene COI (Folmer region) and exhibit different geographical distributions; the H-line displays a Holarctic while the N-line displays a Nearctic distribution with Eastern Canada being the only region of overlap, which suggests signs of divergence. In southwestern Québec, the two lines display evidence of allochronic isolation with N-line larvae appearing in sampled crops almost 2.5 weeks before the H-line (Van der Heyden et al. 2020) and ecological isolation with the N-line being 2.5 times more abundant than the H-line in onions (*Allium*) while the trend is reversed in cruciferous (*Brassica*) crops (Savage et al. 2016). No differences have been detected in either male or female external genitalic structures, and it therefore appears unlikely that mechanical isolation occurs between the two lines. However, as H-line females appear to be selective towards the male with which they copulate, mate discrimination may be a relevant mechanism in pre-mating isolation between the lines (Bush-Beaupré et al. 2022) As COI’s minimum interspecific distance for close relatives in muscoid flies (including *Delia*) is typically between 1.5 and 2.5% (Renaud et al. 2012; Savage et al. 2016), we suspect that the two genetic lines of *D. platura* may in fact be two cryptic species.

As Québec is one of the few Canadian provinces where the distribution of the two *D. platura* lines overlap, crops grown in this area are susceptible to damages caused by both. However, the trends reported by Savage et al. (2016) suggest differences in the potential relative damage contribution by either line depending on the crop. Given that accurate pest identification is crucial for targeted pest control methods, information about the degree to which the H- and N-lines are isolated is needed to develop effective control methods tailored to each line and the crops susceptible to their infestations. Here, we evaluate the reproductive compatibility of the H- and N-lines of *D. platura* with respect to possible pre-mating, post-mating pre-zygotic, and post-zygotic isolation mechanisms using mating probability (insemination probability), pre-oviposition period, egg hatchability as proxies of either pre-mating and/or post-mating pre-zygotic isolation and offspring survival, developmental time, weight, and sex ratio as proxies of post-mating isolation.

## Methods

### Delia platura colonies & experimental stocks

Colonies of the N- and H-lines of *D. platura* were established from wild individuals collected in southern Québec following Bush-Beaupré et al. (2022). Eggs from each main colony were harvested periodically (10 and 9 times for the N- and H-line, respectively) over the course of 10 months and reared on artificial diet containers. Following 16-18 days of development, pupae were harvested, sieved with 1.7-mm mesh to remove very small individuals, and placed in individual plastic vials to be used as adults for the experiment. Voucher specimens were deposited in the Bishop’s University Insect Collection (Sherbrooke, QC, Canada).

### Experimental design

#### Inter- and intra-line crosses

Crosses within and between the two *D. platura* lines were conducted concurrently using a group composition of 30 males to 2 females to maximize female mating probability (Bush-Beaupré et al. 2022. Treatments consisted of intra-line crosses (H-line males with H-line females and N-line males with N-line females), serving as controls, and inter-line crosses (H-line males with N-line females and N-line males with H-line females). Groups were formed of individuals having emerged within 24 hours of each other and placed in a mating arena of approximately 1000 cm^3^ (picture of arena in Supplemental Figure 2). Flies were supplied with distilled water via a dental wick and adult diet *ad libitum* (see Bush-Beaupré et al. 2022 for details). An ovipositional substrate consisting of a 2.0 – 2.5 g piece of rutabaga (*Brassica napus* L.) placed on damp filter paper was supplied and replaced every 2 days for 30 days (see below). Eighteen and 17 replicates of the H-line and N-line intra-line crosses were conducted, respectively, along with 15 replicates for both inter-line crosses.

#### Mating probability

Once oviposition evaluation ceased (see below), all females within a treatment replicate were euthanized (placed in a freezer at -20°C for approximately 24 hours). Females were then stored in 70% ethanol while awaiting dissection. To obtain a measure of the proportion of mated females within the group (mating probability), all three spermathecae of each female within a group were dissected to confirm the presence of sperm masses (Avanesyan et al. 2017). A female was deemed mated if at least one of its spermathecae contained sperm.

#### Oviposition, egg hatchability and pre-oviposition period

Starting from the day on which groups were formed (day 0), oviposition was evaluated every two days by transferring the eggs laid on the ovipositional site to a petri dish with a humid filter paper and counted. As per Bush-Beaupré et al. (2022), we evaluated egg hatchability (# of eggs hatched / # of eggs laid) following 6 days of incubation in a petri dish. Eggs were deemed fertile if they had hatched. In Bush-Beaupré et al. (2022), we determined the pre-oviposition period for both lines to last10 days on average. As such, to allow ample time for mating to occur and obtain representative sample size of egg hatchability, oviposition evaluation ceased after a standardized period of 30 days. Each evaluation day, dead males were replaced with virgin males aged between 1 and 10 days (average number of males replaced for each treatment is shown in Supplemental Figure 1). Dead females were not replaced to avoid bias measurements of the pre-oviposition period. All females survived beyond 10 days for each replicate.

#### Hybrid Development

To evaluate various proxies of progeny developmental success (described below), larvae hatched within 24 hours were transferred as a group of varying density (Table 1) to a plastic container containing approximately 35g of artificial diet (described in Bush-Beaupré et al. 2022. Previous similar experiments had shown 35g to be a generous amount of artificial diet for the development of the number of larvae tested in this experiment (Villeneuve, M.-A., pers. com.). A 2-cm² square was cut out from the lid and covered with fine mesh to allow for air circulation. Starting from the day on which groups were formed (day 0), development was evaluated every 2 days. Evaluation consisted of searching through the artificial diet to retrieve pupae if they were formed. If pupae were found, they were weighed and transferred to individual plastic vials to monitor their development until adult emergence. Evaluation ceased after 30 days for both the larval and pupal stages (total 60 days). After this period, individuals that had not pupated or emerged as adults were considered dead and thus a measure of survival for the larval and pupal stages was obtained. Emerged adults were sexed to measure adult sex ratio.

**Table 1.** Sample sizes to evaluate hybrid *Delia platura* development. Sample sizes for the evaluation of offspring development parameters from crosses between and within the H- and N-line of *Delia platura*. The development of Nm x Hf cross offspring was not evaluated as no offspring were obtained from this cross (see Results section).

| Cross type (male x female line) | Hm x Hf | Hm x Nf | Nm x Nf |
| --- | --- | --- | --- |
| Number of crosses sampled | 12 | 12 | 13 |
| Mean larvae #/cross | 20 | 16 | 21 |
| Min & max larvae #/cross | 9 – 30 | 8 – 30 | 9 – 30 |
| Total larvae # evaluated (developmental time & survival) | 243 | 197 | 280 |
| Total pupae # weighed | 120 | 140 | 163 |

#### Statistical analyses

Unless otherwise indicated, model fitting was done using the glmmTMB package (v.1.1.2; Brooks et al., 2017) and model diagnostics were inspected using the DHARMA package (v.0.4.3; Hartig, 2021) within the R environment (R Core Team 2021).

#### Mating probability

Female mating probability was modeled using a Generalized Linear Model with a binomial error distribution and complementary log-log (cloglog) link function. Cross type (Hm x Hf, Nm x Nf, Hm x Nf, Nm x Hf (male line x female line)) was used as a covariate with the Hm x Nf cross as the baseline group. Female and male lot ID were initially included in the model as random effects to control for the variance in mating probability between lots. However, these variables had a small variance (1.421e^-11^ and 8.406e^-10^ for female and male lot ID, respectively) and hindered model fit. These two variables were thus removed from the model. As some females did not survive for the entire 30-day oviposition evaluation period, female mating probability was partly influenced by lifespan. To take this into account, we counted the number of female-day (number of females still alive per evaluation day) and included this in the model as an offset term (log(total female-days)).

#### Egg hatchability

The proportion of eggs hatched from the number of eggs laid was modeled using a Generalized Linear Mixed Model with a binomial error distribution and complementary log-log (cloglog) link function. As only one female in the Nm x Hf cross mated and laid fertile eggs (see results section), only the three other types of crosses were included as covariates with the Hm x Nf cross as the baseline group. In this model, female and male lot ID were found to have a substantial variance and improved model fit and were thus included as random effects. Following the same logic as in the female mating probability model, log(total female-days) was included as an offset term.

#### Pre-oviposition period

The time (in days) until the first fertile eggs were laid in each group was modeled using a Generalized Linear Mixed Model with a gamma error distribution and log link function. The Hm x Nf cross was compared to the Nm x Nf cross to evaluate differences in the time N-line females took to accept either hetero- or con -specific males. Here, we assume that the duration of mating and the delay between insemination and oviposition is the same for both cross types. As only one H-line female mated with an N-line male (see Results), the pre-oviposition period of the Hm x Hf and Nm x Hf crosses were not compared statistically. Male and female lot ID were included as random effects.

#### Hybrid survival

The proportion of individuals that survived from the larval stage to the pupal stage was modeled using a Generalized Linear Mixed Model with a binomial error distribution and logit link. The three cross types that produced larvae were included as covariates with the Hm x Nf cross as the baseline group. As individuals that developed within a single artificial diet container are not fully independent, a group ID random effect was thus included in the model along with female and male lot ID. Although the variance of each of these variables was low, the combination of the three random variables showed an improved model fit and were thus retained. Survival from the pupal to the adult stage and from the larval to the adult stage was modeled in the same manner (3 models total).

#### Hybrid developmental time

A single Bayesian Generalized Linear Mixed Model was used to model developmental time (in days) from the larval to pupal stages and from pupal stage to adult emergence. Both the mean and shape (degree of dispersion) of the gamma distribution were modeled. We modeled the shape parameter as a means to describe the degree to which developmental time may vary among individuals within groups between hybrids and the parental lines. We hypothesized that genetic incompatibilities may result in developmental times being more highly dispersed (i.e., some individuals may have shorter/longer developmental times compared to the average). In the Gamma distribution, this is achieved by modelling treatment effects on the shape parameter; a higher shape parameter value translates to a lower dispersion and variance in development time. The three cross types were included as covariates with the Hm x Nf cross as the baseline group along with sex (female or male), developmental stage (larval or pupal) and their interaction for both the mean and shape parameters. Individual ID was included as a random effect for the mean parameter to control variation in developmental time between the larval and pupal stage for each individual along with a random slope variable for developmental stage and group ID. The random slope variable for developmental stage and group ID was also included for the shape parameter. Additionally, to facilitate the estimation of the random effect structure of the shape parameter, a variance-covariance matrix was included between the mean and shape random slope variables. As such, a positive correlation between the mean and shape random slopes can be understood as a decreased dispersion as the mean increases. Complete model equations and priors are included in Supplemental Figure 3. Model fitting was performed with the package brms (v.2.16.3; Burkner, 2017) in R.

#### Pupal weight

Pupal weight in milligrams was modeled using a Linear Mixed Model with a Gaussian error distribution and identity link. The three cross types were included as covariates with the Hm x Nf cross as the baseline group along with sex (female or male) and their interaction. In this model, only group ID was included as a random effect due to model convergence issues when including female and male lot ID.

#### Adult sex ratio

The proportion of male to female adults was modeled using a Generalized Linear Mixed Model with a binomial error distribution and logit link. The three cross types were included as covariates with the Hm x Nf cross as the baseline group along with sex (female or male) and their interaction. In this model, group ID, female and male lot ID were included as random variables.

### Results

#### Mating probability

The overall mean proportion of mated females was 51.5% ± 37.5% (SD). Out of 15 replica (30 females), a single H-line female mated with an N-line male, representing only 3% of mated females of that cross type whereas the Hm x Nf, Hm x Hf and Nm x Nf crosses all displayed proportions of mated female equal to or higher than 65% (i.e., 65, 65 and 75 %, respectively; Figure 1A, Supplemental Table 1).

**Figure 1.**
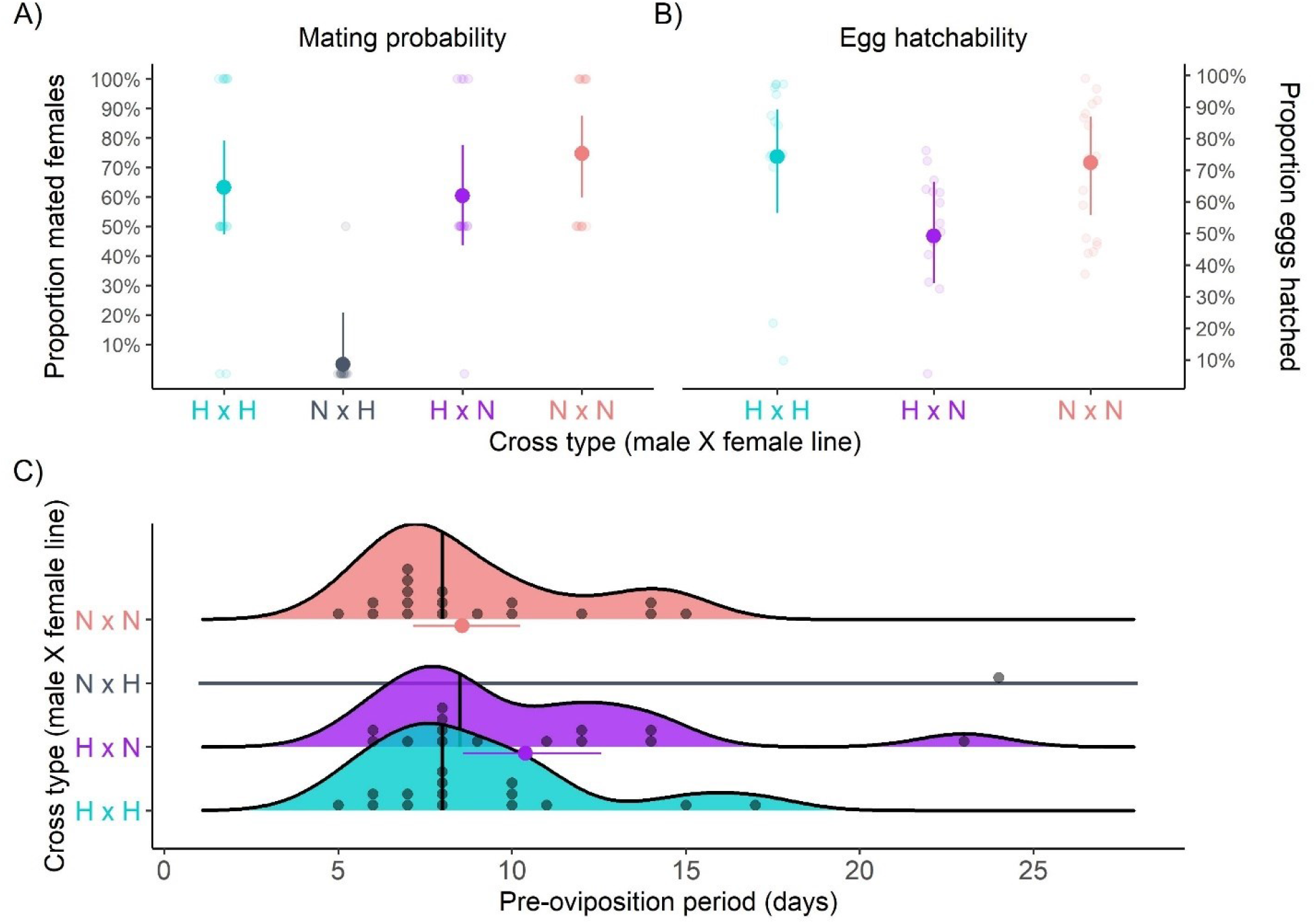
Effect of *Delia platura* cross type on mating success, egg hatchability and pre-oviposition period. Model predictions of the effect of *Delia platura* cross type on mating probability (A) and egg hatchability (B) error bars display 95% confidence intervals. (C) displays the raw data distributions of the first day on which fertile eggs were laid (pre-oviposition period) for each *D. platura* cross type along with model-predicted means and 95% confidence intervals for the pre-oviposition period between Nm x Nf and Hm x Nf crosses.

#### Egg hatchability

The overall mean proportion of eggs hatched was 65.3% ± 24.8%. The proportion of hatched eggs laid by N-line females mated with H-line males was 50% compared to 75% in the intra-line crosses (Figure 1B, Supplemental Table 1). The only mated H-line female in the Nm x Hf cross laid 59 fertile eggs out of a total of 477 eggs laid (12.4% hatchability).

#### Pre-oviposition period

The overall mean pre-oviposition period for both Hm x Nf and Nm x Nf crosses was 9.7 ± 4.1 days. These two cross types had comparable predicted mean pre-oviposition (10.4 and 8.6 days for the Hm x Nf and Nm x Nf crosses, respectively; Figure 1C, Supplemental Table 1). The Hm x Hf cross had a mean pre-oviposition period of 9.1 ± 3.3 days while the only female from the Nm x Hf cross to lay fertile eggs did so after 24 days.

#### Hybrid survival

The overall mean survival from the larval to pupal stage was 87.5% ± 33.1%. Hm x Nf hybrids had a similar survival compared to the parental lines (∼95%; Figure 2, Supplemental Table 2). The overall mean survival from the pupal to adult stage for the larvae that survived to the pupal stage was 90.9% ± 28.8%. Hm x Nf hybrid survival was comparable to the H-line and was 5% higher compared to the N-line (95% vs 90%; Figure 2, Supplemental Table 2). The overall mean survival from the larval to adult stage was 79.5% ± 40.4%. Again, Hm x Nf hybrid survival was comparable to that of the intra H-line crosses and was nearly 20% higher compared to that of the intra N-line crosses (∼92% vs 75%; Figure 2, Supplemental Table 2).

**Figure 2.**
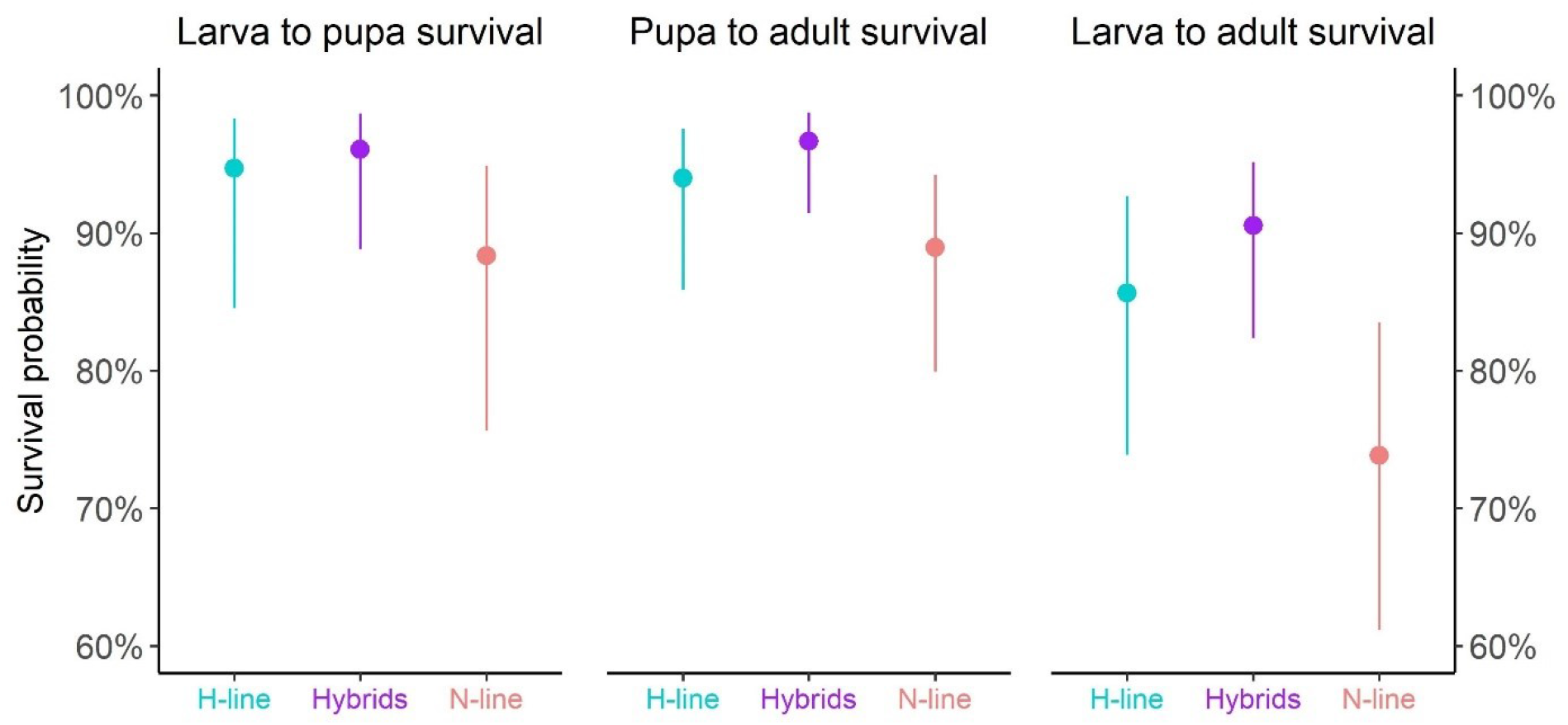
Effect of *Delia platura* cross type on offspring survival. Model predictions for the effect of *Delia platura* parental and Hm x Nf hybrid lines on survival probability from the larval to pupal, pupal to adult and larval to adult life stages. Error bars depict 95% confidence intervals.

#### Hybrid developmental time

The overall mean developmental time from the larval to pupal stage was 11.5 ± 2.6 days whereas developmental time from the pupal to adult emergence stage was 11.8 ± 2.6 days. Mean developmental time was similar between the hybrids and parental lines throughout both developmental stages and between sexes. The shape parameter (dispersion) was also similar between the hybrids and parental lines throughout both developmental stages and between sexes (Figure 3; Supplemental Figure 4). The hybrids were thus equivalently variable in developmental time compared to the parental lines.

**Figure 3.**
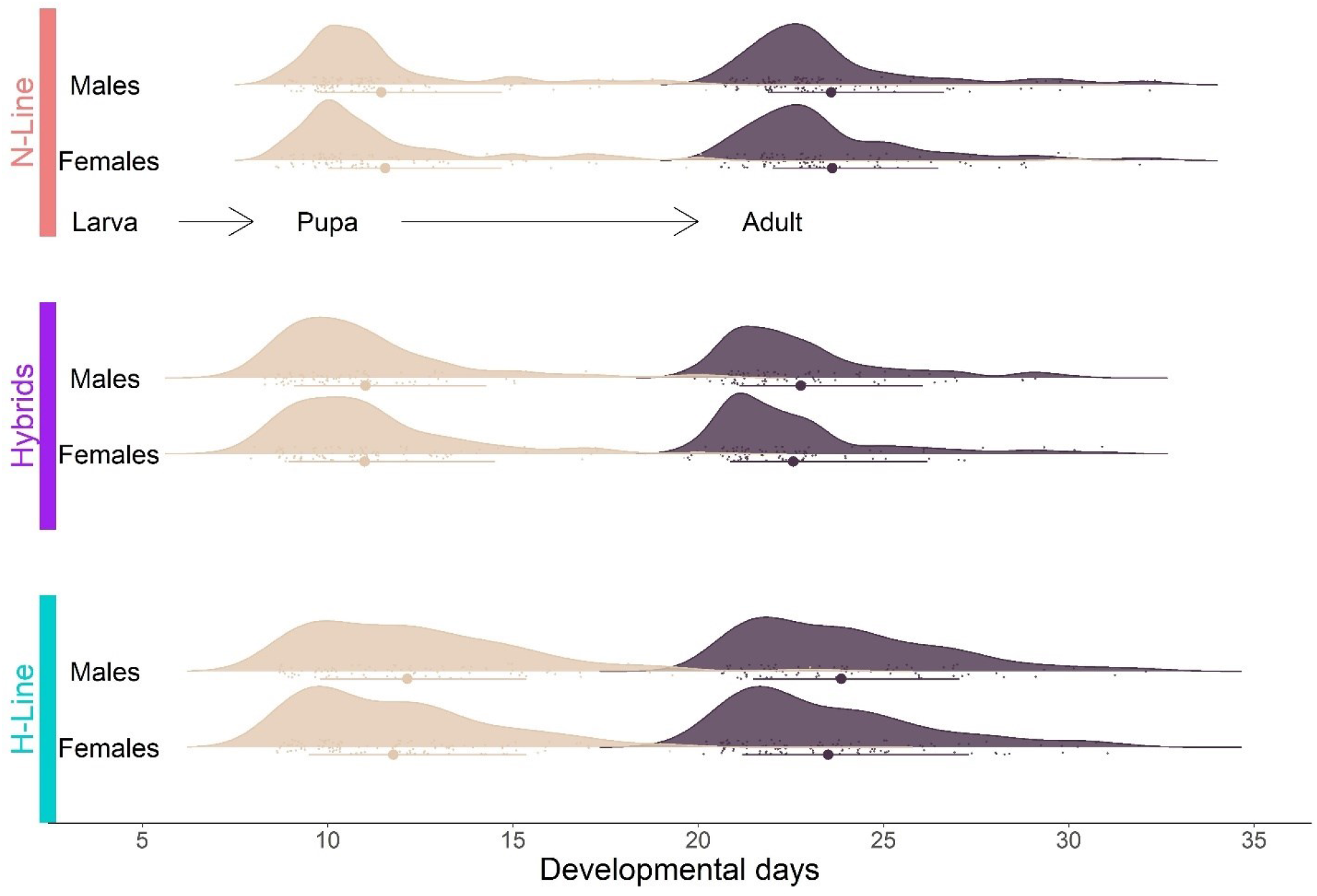
Effect of *Delia platura* cross type on offspring developmental time. Model predictions for the effect of *Delia platura* parental and Hm x Nf hybrid lines and sex on developmental time from the larval to pupal and pupal to adult life stages. Error bars depict 95% confidence intervals.

#### Pupal weight

The overall mean pupal weight was 8.8 mg ± 1.3 mg. Males weighed approximately 1 mg more than females for the Hm x Nf hybrids and parental lines. On average, hybrid pupal weight was comparable to N-line pupae and 15 mg more than H-line pupae (Figure 4A, Supplemental Table 3).

**Figure 4.**
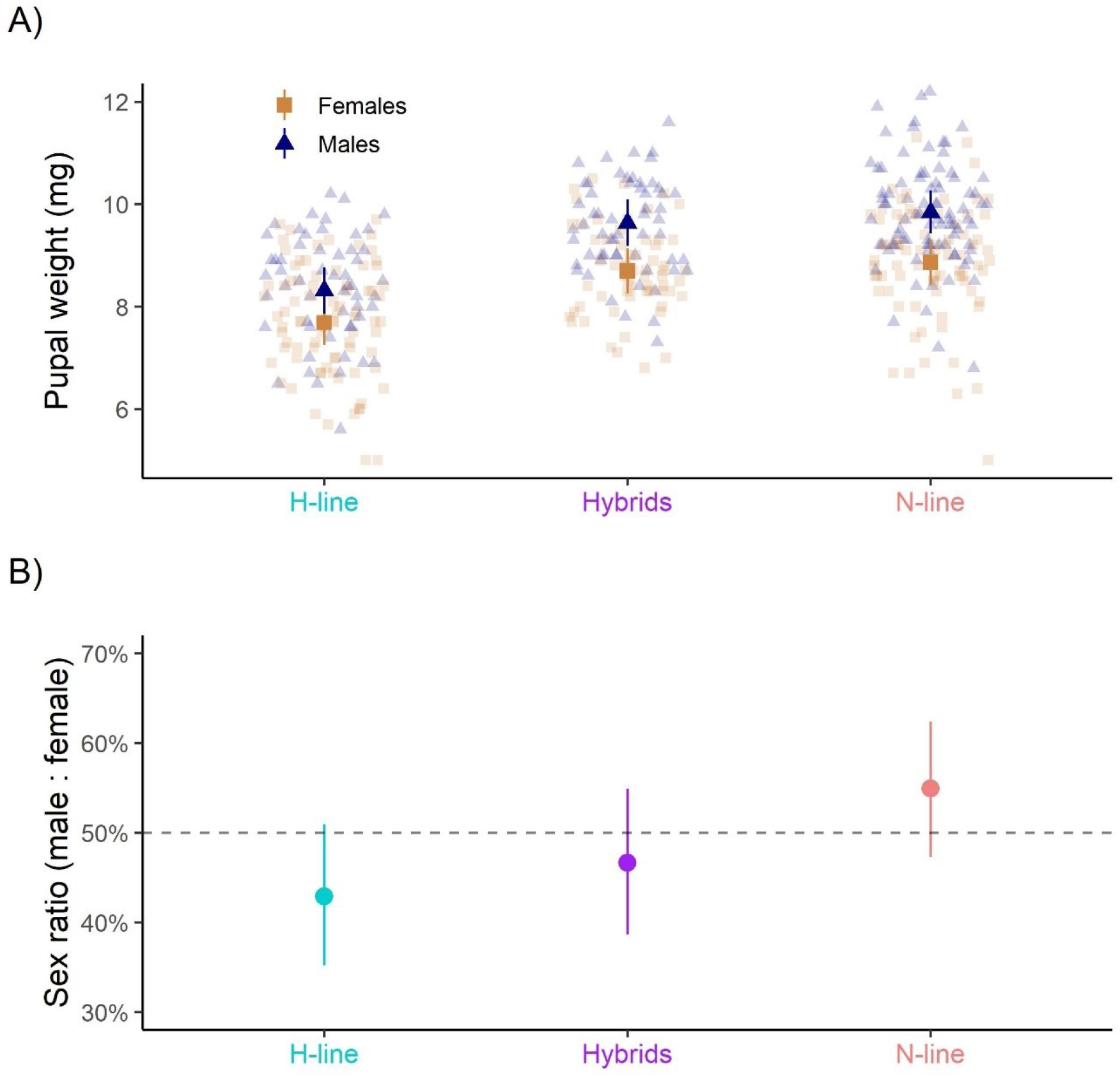
Effect of *Delia platura* cross type on offspring pupal weight and sex ratio. Model predictions for the effect of *Delia platura* parental and Hm x Nf hybrid lines and sex on pupal weight (A) and the effect of parental and hybrid lines on offspring sex ratio (B). Error bars depict 95% confidence intervals.

#### Adult sex ratio

The overall mean male proportion was 48.3% ± 50.0%. Neither the Hm x Nf hybrids nor parental lines deviated substantially from this mean (predicted means of 46.7%, 54.9% and 42.9% for the Hm x Nf hybrids, pure N-line and pure H-line, respectively; Figure 4B, Supplemental Table 3).

## Discussion

The reproductive compatibility between the N- and H-line of *D. platura* was investigated by evaluating multiple proxies of pre-mating, post-mating pre-zygotic, and post-zygotic isolation. We uncovered asymmetrical pre-mating isolation between the lines, with only one H-line female out of thirty being inseminated byan N-line male (and laying approximately 12% fertile eggs) while N-line females readily mated with H-line males. However, the eggs laid by N-line females in the inter-line crosses hatched at a 25% lower rate than the intra-line crosses suggesting either post-mating pre-zygotic or post-zygotic isolation. The larvae that successfully hatched from this inter-line cross had a high degree of developmental success according to the proxies evaluated: survival to adult emergence from puparia, larval and pupal developmental time, pupal weight, and adult sex ratio. Thus, we uncovered no evidence of post-zygotic isolation in the development of hybrids resulting from Hm x Nf cross. Our results point to partial reproductive isolation between the two lines, suggesting that they are in fact two unique biological entities which could in turn influence the development of species-specific control methods for either line.

### Mating probability

Pre-mating isolation typically impedes mating success (i.e., insemination; Coyne & Orr, 2004). If the H- and N-line of *D. platura* were entirely isolated due to pre-mating barriers, we would expect no sperm transfer to occur in all inter-line crosses. Our results, however, point to a partial asymmetrical reproductive barrier between the two lines. A possible isolating mechanism explaining the very low mating success for Nm x Hf crosses may be in the form of behavioral pre-mating isolation. Mate recognition and discrimination during courtship behavior is known to negatively impact the mating success of inter-genetic crosses between many species/biotypes (Elbaz et al. 2010; Rull et al. 2013; Boumans and Johnsen 2014). It has been previously shown that H-line females are choosier towards the male with which they mate compared to the N-line (Bush-Beaupré et al. 2022) and this high degree of mating selectivity may point to an increase potential for the H-line females to discriminate between males of either line. However, as emphasized by Fraser & Boake (1997), behavioral isolation through male or female choice should not be inferred solely by mating success data but rather supplemented with detailed observations and experiments pertaining to the mechanisms of mate choice.

Although behavioral pre-reproductive isolation may be present between the two lines, other pre-zygotic barriers may also impede Nm x Hf crosses. The observed variable used as a proxy for mating in this study was the presence of sperm in the spermathecae. It is, however, possible for copulation to have occurred in this type of cross without sperm transfer. For example, the behavior of either female or male during copulation may not have been appropriate and thus led to a failure in sperm transfer, a phenomenon documented in inter-specific crosses of *Nasonia* (Hymenoptera: Pteromalidae) wasps (Buellesbach et al. 2014). Mechanical isolation through differences in internal genital morphology may also contribute to pre-reproductive isolation between the lines (Masly 2012). The relative strengths of potential pre-zygotic isolating mechanisms remain to be uncovered.

### Pre-oviposition period

As the pre-oviposition period of N-line females mated with H-line males was similar to that of N-line females involved in intra-line crosses, it appears that N-line females do not exhibit resistance towards mating with H-line males, further displaying the lack of pre-reproductive barrier in this type of cross. In contrast, the single H-line female mated with an N-line male took more than twice the average time observed in all other cross types. Based on the assumption that *D. platura* only mates once, as documented for its close relative, the onion maggot (*Delia antiqua*) (as *Hylemya antiqua* in Martin & McEwen, 1982), the long pre-oviposition period observed in that H-line female suggests that it was reluctant to mate with an N-line male or use its sperm to fertilize her eggs. While the two aforementioned processes are not mutually exclusive, it is possible that our observed trends are similar to trends that have been observed in crosses between the B and Q biotypes of *Bemisia tabaci* (Hemiptera: Alerodidae) where biotype B females spent a shorter amount of time being courted by males in intra-specific crosses compared to inter-specific crosses, whereas biotype Q females had negligeable differences in time spent being courted in either cross type (Elbaz et al. 2010). As both intra-line crosses of *D. platura* had similar pre-oviposition periods, it can be inferred that the age of sexual maturity is approximately the same for of both lines and that allochronic isolation due to sexual maturity is therefore not an isolation mechanism between the two lines.

### Egg hatchability

Post-mating pre-zygotic isolation in the form of reduced egg hatchability is commonly observed among insect taxa (Price et al. 2001; Matsubayashi and Katakura 2009; Marshall et al. 2009; Garlovsky and Snook 2018; Hill et al. 2021). In our study, the eggs laid by N-line females mated with H-line males hatched at a 25% lower proportion compared to the intra-line crosses, suggesting a reduction in egg fertilization, and thus zygote formation. While post-mating pre-zygotic isolation is still a developing area of research, many studies have demonstrated the involvement of seminal fluid proteins in egg fertilization failure and defective sperm storage (Marshall et al. 2009; Larson et al. 2012; Ahmed-Braimah 2016; Garlovsky et al. 2020; Hill et al. 2021). A plausible explanation for our results may be differences in seminal fluid proteins between males of both lines which could affect egg fertilization rates and/or female sperm utilization. Incompatibilities in internal genital morphology, impeding the ability for sperm to reach eggs, between the lines may also have led to zygote formation failure (Masly 2012). However, as we did not dissect the unhatched eggs to evaluate the presence of unhatched larvae, it is also possible that the low hatching proportion of eggs laid by N-line females mated with H-line males is due to failure in embryonic development. Thus, it is also likely that the observed form of isolation is post-zygotic rather than post-mating pre-zygotic. Although zygote formation and/or embryonic development was reduced in our Nm x Hf crosses, approximately 50% of the eggs laid produced viable larvae and thus post-mating pre-zygotic or post-zygotic isolation is only partially present in crosses between N-line females and H-line males.

### Hybrids

Zygote formation does not imply complete reproductive compatibility between populations. Indeed, post-zygotic isolation mechanisms in the form of either hybrid unviability or suppression (selection against hybrids) are often observed (dos Santos et al. 2001; Naisbit et al. 2001; Coyne and Orr 2004; Forister 2005; Zhao et al. 2005; Gibson et al. 2013; Phadnis et al. 2015; Roriz et al. 2017). Many proxies, such as the ones evaluated in our study, may be used to assess hybrid viability including hybrid survival, developmental time, pupal weight, and sex ratio (Nakakita and Imura 1981; Laurie 1997; Ording et al. 2010; Roriz et al. 2017). If post-zygotic isolation in the form of hybrid unviability were to be present between the H-line males and N-line females of *D. platura*, we would expect differences between the hybrids and the parental lines for the measured proxies.. However, none of the aforementioned proxies measured suggest a form of post-zygotic isolation through hybrid unviability; hybrids performed at the same rate as their parents in terms of survival (similar to H-line), developmental time (similar mean and variability as both lines), pupal weight (similar to N-line), and sex ratio (similar to both lines). The evidence obtained here point to successful hybrid development from the larval stage to adult emergence similar to trends obtained among multiple insect taxa (Bierbaum and Bush 1990; Velásquez-Vélez et al. 2011; Petrucco-Toffolo et al. 2018; He et al. 2021). It is, however, noteworthy that the results presented here pertain solely to the development of the F1 hybrid generation and, as such, additional data on their longevity and fertility will be necessary to fully assess hybrid viability.

In many cases, insect hybridization may lead to range expansion, increase resistance to management techniques, along with outbreaks as host range increases (Anderson et al., 2018; Andersen et al., 2019; Corrêa et al., 2019). It is thus important to monitor these possible hybrid populations to better understand the possible outcomes of such hybridization. Considering the hybridization potential between H-line males and N-line females of *Delia platura* in laboratory conditions, genetic sampling of field populations in the area of H- and N-line distribution overlap (eastern North America) is required to determine the population genetic structure and potentially uncover hybrid populations. To uncover probable hybrid populations, data on additional genetic markers, preferably nuclear rather than mitochondrial markers, is needed. Data on these markers would help to determine if the genetic differences between the lines go beyond CO1 haplotypes. If such populations exist, they may exhibit different phenotypic trends in term of their phenology, distribution, and host range thus affecting the methods by which growers may control the damage caused in their fields

Our results are congruent with findings obtained by (Jennings et al. 2014) with laboratory-based crosses involving three allopatric populations of *Drosophila montana*. In their study system, females from the Vancouver population showed clear preference towards males of their own population compared to males from Colorado and Oulanka populations in mate-choice trials, suggesting asymmetric pre-mating isolation as we observed in crosses between H-line females and N-line males of *Delia platura*. Additionally, in some of the inter-population crosses where mating readily occurred, the authors observed reduced egg hatchability and, after dissecting female reproductive tracts, determined that sperm had been successfully transferred, stored and was motile. However, dissection of unhatched eggs revealed that a large proportion of these eggs had been unfertilized, suggesting incompatibilities in surface proteins between sperm and egg. Analogous to our findings of reduced egg hatchability in crosses between H-line males and N-line females, this post-mating pre-zygotic barrier was incomplete as 30 to 60% of the eggs produced in these crosses were successfully fertilized and developed. Although we did not evaluate egg fertilization success, our results point to a deficiency in fertilization rates or survival failure in the early stages of larval development. The authors observed no post-zygotic isolation in the crosses that successfully produced progeny as the hybrids had a high rate of survival and produced F2 progeny. While we did not evaluate hybrid fertility in our study, we did observe high hybrid development success from the larval hatching to adult emergence stages. As Jennings et al. (2014) point out, pre-mating and post-mating barriers can develop independently from one another and so, it is likely that the hybrids produced by H-line males and N-line females are fertile and produce F2 progeny. The results obtained by Jennings et al. (2014) further emphasize the need to evaluate *D. platura* hybrid persistence.

### Two biological entities

As emphasized by Carstens et al. (2013), species delimitation analyses are strongest when assessing both genetic and biological lines of evidence and their congruence. A proper understanding of the speciation process in insects relies on a study of the evolutionary processes that lead to the evolution of speciation phenotypes, their genetic architecture and how they influence patterns of gene flow along with the relationship between their divergence and species boundaries (Mullen & Shaw, 2014). One probable speciation phenotype within the *D. platura* H- and N-line complex is the apparent mate selectivity that H-line females exhibit as determined in Bush-Beaupré et al. (2022) and emphasized in our study. However, we have yet to uncover the processes that led to the evolution of such phenotype and, more importantly, if this phenotype (among others) is present in field conditions. While the evolutionary and taxonomic status of the two *D. platura* lines may not, at present, be determined, the differences in the mating system of the two lines (Bush-Beaupré et al. 2022) along with the pre-mating and pre-zygotic barriers exposed in this study suggest that the two lines are two biological entities. Due to these biological differences, we propose the provisional designation of the term biotype for the H- and N-lines of *D. platura* as suggested by Diehl & Bush (1984) until their evolutionary and taxonomic status be resolved. Such designation may serve to emphasize the biological differences between the biotypes among the agronomic community and hopefully guide future research on control methods tailored to each biotype.

### Future Studies

Our results suggest various avenues of future studies on the reproductive isolation of the *D. platura* H- and N-lines. Although our results suggest behavioral pre-zygotic isolation between H-line females and N-line males, future work should investigate the presence of other isolating mechanisms and their relative strengths. Additionally, further research in male seminal fluid composition, female sperm utilization, internal genital morphology, and embryonic development is needed to uncover the mechanisms underlying either the post-mating pre-zygotic or post-zygotic isolation observed between H-line males and N-line females. Finally, hybrid adult survival and fertility should be evaluated in future experiments to confirm *D. platura* hybrid viability across their entire lifecycle and future generations.

### Applications

Targeted pest control methods rely on proper identification of the identity of the pest involved. One such method is the sterile insect technique (SIT) that is rendered inefficient if released sterile individuals do not mate with the targeted population. Genetically distinct populations may differ in some of their biological traits and occurrence in the field but are known to readily interbreed given a particular setting (Pereira et al. 2007; Ahmad et al. 2018). In these cases, mass rearing and release of only one of the biotypes could potentially be effective for the control of both. However, biotypes of a given species may interbreed but in an asymmetrical pattern, reducing the effectiveness of SIT if only one of the strains is released (Roriz et al. 2017). Given that the H- and N-biotypes of *D. platura* are asymmetrically reproductively compatible, we suggest the release of individuals of the same biotype as the targeted population if such method should be employed. Other biological control agents such as entomopathogenic fungi, parasitoids, and nematodes may have small host ranges and sometimes may be host-specific (Davidson and Chandler 2005; Jensen et al. 2006; Eilenberg and Jensen 2018). Recognition of the two *D. platura* biotypes as separate biological entities may allow future research to focus on determining the control agents specific to each biotype. For example, when exposed to seven species of nematodes, *D. platura* in Colombia (presumably N-line) was most susceptible to *Steinernema* sp.3 (Jaramillo et al. 2013). It is likely, however, that the species of nematode (of other biological control agent) found to be effective against biotype N is not as effective in controlling biotype H populations. We therefore recommend testing and developing control methods targeted to a specific *D. platura* biotype as the efficacy of a method may not be equivalent for both biotypes.

## Acknowledgements

We are grateful to Marc-André Villeneuve and the students at Collège Montmorency for logistical support. This work was funded by Agriculture and Agri-Food Canada, The Fruit & Vegetable Growers of Canada (Canadian AgriScience Cluster for Horticulture program) and Bishop’s University.

