## Supplementary material for "REPRODUCTIVE COMPATIBILITY OF TWO LINES OF *DELIA PLATURA* (DIPTERA: ANTHOMYIIDAE)"

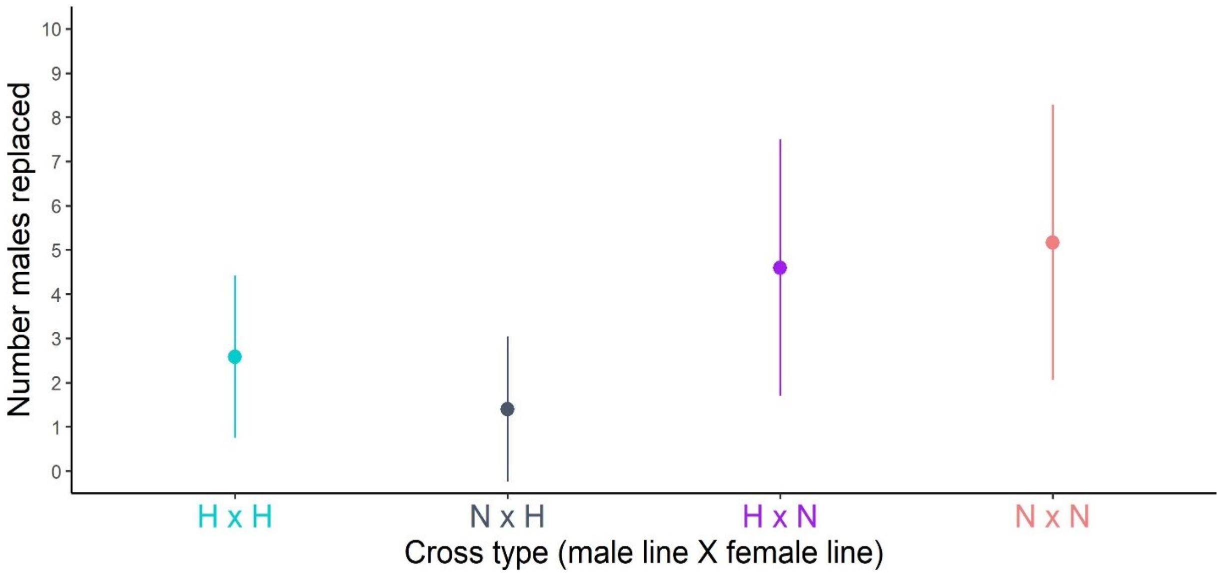


**Supplemental Figure 1.** **Average (± SD) number of dead males replaced for each *Delia platura* cross type**

Average (± SD) number of dead males replaced for each treatment evaluating the effect of cross type on mating probability, pre-oviposition period and egg hatchability for the H- and N- lines of *D. platura*.

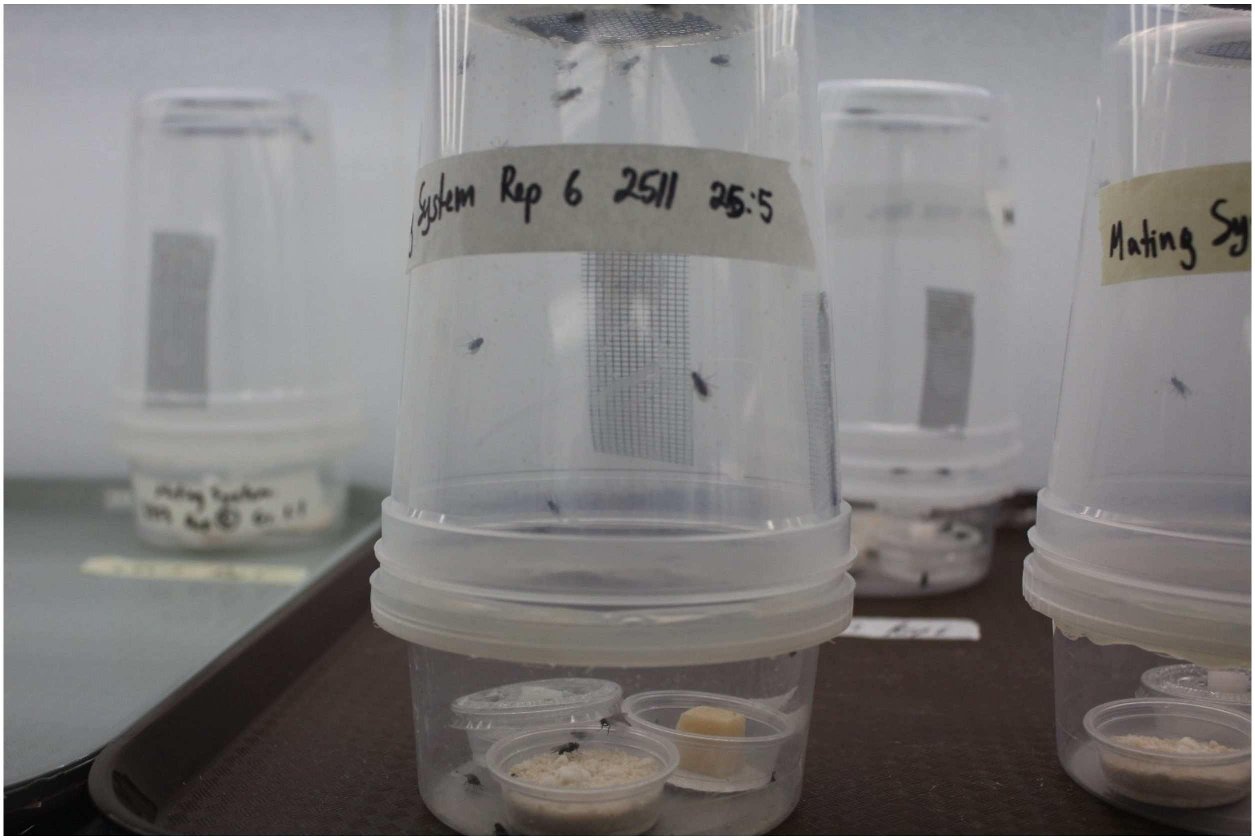


**Supplemental Figure 2.** **Arena used for the reproductive compatibility experiment**

Arena used to evaluate the effect of cross type on mating probability, pre-oviposition period and egg hatchability for the H- and N- lines of *D. platura*.

**Supplemental Table 1.** **Output of statistical analyses for the effect of *Delia platura* cross type on mating probability, egg hatchability and pre-oviposition period**

| | **Mating probability** | | **Egg hatchability** | | **Pre-oviposition period** | |
|---|---|---|---|---|---|---|
| *Predictors* | *Estimate* | *CI (95%)* | *Estimate* | *CI (95%)* | *Estimate* | *CI (95%)* |
| (Intercept) | -4.15 | -4.63 – -3.67 | -4.45 | -4.93 – -3.98 | 2.34 | 2.15 – 2.53 |
| cross [nxh] | -3.34 | -5.36 – -1.33 | | | | |
| cross [hxh] | 0.08 | -0.58 – 0.73 | 0.70 | 0.17 – 1.22 | | |
| cross [nxn] | 0.40 | -0.23 – 1.03 | 0.64 | 0.19 – 1.09 | -0.19 | -0.42 – 0.04 |
| **Random Effects** | | | | | | |
| $\sigma^2$ | | | $0.33_{\text{female.lot.ID}}$ | | $0.00_{\text{male.lot.ID}}$ | |
| | | | $0.22_{\text{male.lot.ID}}$ | | $0.02_{\text{female.lot.ID}}$ | |
| $R^2$ conditional / $R^2$ marginal | NA / 0.577 | | 0.042 / 0.285 | | 0.092 / NA | |

Estimates and corresponding 95% confidence intervals (CI) of the effects of *D. platura* cross type on mating probability, egg hatchability, and pre-oviposition period from Generalized Linear Mixed Models. A binomial error distribution with a cloglog link function was used for the mating probability and egg hatchability analyses while a gamma distribution with a log link function was used for the pre-oviposition period analysis. $\sigma^2$ represents the variance of the random effects (in subscript next to value). Note that the estimates for the mating probability and egg hatchability analyses are on the cloglog scale while the estimates are on the log scale for the pre-oviposition period analysis.

**Supplemental Table 2. Output of statistical analyses for the effect of D*elia platura* cross type on offspring survival probability**

| | Larva to pupa survival | | Pupa to adult survival | | Larva to adult survival | |
|---|---|---|---|---|---|---|
| *Predictors* | *Estimate* | *CI (95%)* | *Estimate* | *CI (95%)* | *Estimate* | *CI (95%)* |
| (Intercept) | 3.19 | 2.07 – 4.32 | 3.37 | 2.37 – 4.38 | 2.26 | 1.54 – 2.98 |
| cross [nxn] | -1.17 | -2.38 – 0.04 | -1.29 | -2.48 – -0.09 | -1.22 | -1.98 – -0.46 |
| cross [hxh] | -0.31 | -1.86 – 1.23 | -0.62 | -1.90 – 0.65 | -0.47 | -1.49 – 0.55 |
| **Random Effects** | | | | | | |
| $\sigma^2$ | $0.80_{\text{group.ID}}$ | | $0.40_{\text{group.ID}}$ | | $0.23_{\text{group.ID}}$ | |
| | $0.95_{\text{female.lot.ID}}$ | | $0.00_{\text{female.lot.ID}}$ | | $0.49_{\text{female.lot.ID}}$ | |
| | $0.00_{\text{male.lot.ID}}$ | | $0.38_{\text{male.lot.ID}}$ | | $0.00_{\text{male.lot.ID}}$ | |
| Marginal $R^2$ / Conditional $R^2$ | 0.071 / NA | | 0.078 / NA | | 0.072 / NA | |

Estimates and corresponding 95% confidence intervals (CI) of the effects of *D. platura* cross type on offspring survival from the larval to pupal, pupal to adult and larval to adult stages from Generalized Linear Mixed Models. A binomial error distribution with a logit link function was used for all analyses. $\sigma^2$ represents the variance of the random effects (in subscript next to value). Note that the estimates are on the logit scale.

Developmental time ~ Gamma(μ, α) where μ = mean, α = shape

Log(μ) ~ cross type (H-line/N-line/Hybrids) * developmental stage (larval/pupal) * sex (female/male) + (1|larva.ID) + (1 + developmental stage|g|group.ID)

Log(α) ~ cross type (H-line/N-line/Hybrids) * developmental stage (larval/pupal) * sex (female/male) + (1 + developmental stage|g|group.ID)

Where g = 4 dimensional variance-covariance matrix

Priors:

μ ~ Normal(2.5, 1)

fixed effects for μ & α ~ Normal(0, 0.5)

α ~ Normal(3, 1)

larva.ID ~ Cauchy(0, 3)

group.ID ~ Cauchy(0, 3)

g ~ LKJ(1)

**Supplemental Figure 3. Model equation and priors for Bayesian Generalized Linear Mixed Model used for evaluating the effect of *Delia platura* cross type, developmental stage and sex on progeny developmental time**

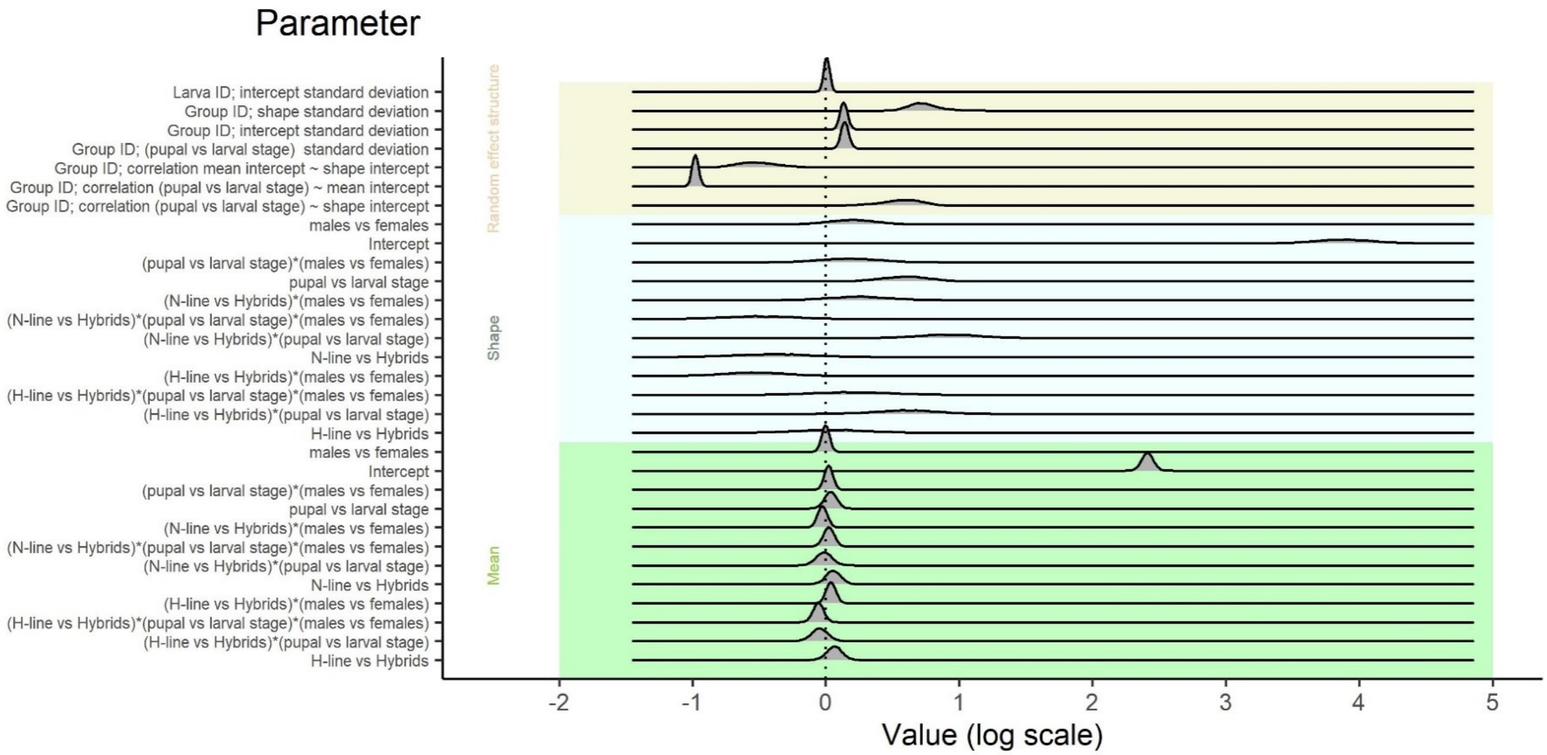


**Supplemental Figure 4.** **Posterior distribution of parameters estimated for the effect of *D. platura* cross type, developmental stage and sex on progeny developmental time obtained from a Bayesian Gamma Generalized Linear Mixed Model**

**Supplemental Table 3. Output of statistical analyses for the effect of D*elia platura* cross type on offspring pupal weight and sex ratio**

| | Pupal weight | | Offspring sex ratio | |
|---|---|---|---|---|
| *Predictors* | *Estimate* | *CI (95%)* | *Estimate* | *CI (95%)* |
| (Intercept) | 8.69 | 8.25 – 9.14 | -0.13 | -0.46 – 0.20 |
| cross [nxn] | 0.17 | -0.46 – 0.79 | 0.33 | -0.13 – 0.79 |
| cross [hxh] | -1.01 | -1.63 – -0.39 | -0.15 | -0.60 – 0.30 |
| sex [male] | 0.94 | 0.59 – 1.29 | | |
| cross [nxn] * sex [male] | 0.04 | -0.42 – 0.50 | | |
| cross [hxh * sex [male] | -0.32 | -0.79 – 0.16 | | |
| **Random Effects** | | | | |
| $\sigma^2$ | 0.32 $_{\text{group.ID}}$ | | 0.02 $_{\text{group.ID}}$ | |
| | | | 0.00 $_{\text{female.lot.ID}}$ | |
| | | | 0.02 $_{\text{male.lot.ID}}$ | |
| Marginal $R^2$ / Conditional $R^2$ | 0.333 / 0.514 | | 0.013 / NA | |

Estimates and corresponding 95% confidence intervals (CI) of the effects of *D. platura* cross type on offspring pupal weight and sex ratio from Generalized Linear Mixed Models. A gaussian error distribution with an identity link function was used for the pupal weight analysis while a binomial distribution with a logit link function was used for the offspring sex ratio analysis. $\sigma^2$ represents the variance of the random effects (in subscript next to value). Note that the estimates of the pupal weight analysis are on the response scale while the estimates are on the logit scale for the offspring sex ratio analysis.
